# Genetic Diversity and Antifungal Susceptibility among Ecological Niche Populations of *Aspergillus flavus* in Yaoundé, Cameroon

**DOI:** 10.64898/2026.07.22.740195

**Authors:** Marius P. Ngouanom Kuate, Kesini Ramkumar, Jezreel Dalmieda, Javeriya Nadeem, Javid Hemmati, Celine N. Nkenfou, Roland N. Ndip, Jianping Xu

## Abstract

*Aspergillus flavus* is a ubiquitous fungus commonly found in a variety of ecological niches including soil, crops, the air, and humans. In humans, it is the second leading cause of invasive aspergillosis (IA) and is linked to other illnesses through contamination of agricultural products with aflatoxins. As such, *A. flavus* threatens food safety, economic wellbeing, and human health, particularly in less developed regions and countries such as Cameroon. To mitigate these effects, farmers and clinicians rely on triazoles to reduce aflatoxin contamination and treat IA. However, a consequence of increasing triazole use is the emergence and spread of triazole resistance in both agriculture and clinics. To identify the prevalence of triazole resistance and the potential relationships among triazole resistant and susceptible strains, this study investigated antifungal susceptibility to both clinical and agricultural triazoles and analyzed genetic variation among various ecological niche populations of *A. flavus* in Cameroon. Strain genotypes were obtained through analysis of six polymorphic microsatellite markers. Our analyses revealed that 30.3% (17/56) of the strains were resistant to at least one of the four triazoles, with increased minimum inhibitory concentrations found among crop-isolated strains. A Permutational Multivariate Analysis of Variance suggested limited ecological niche-based clustering of genotypes, consistent with frequent gene flow and dispersal among ecological niches. A multilocus linkage disequilibrium analysis revealed evidence of non-random recombination in *A. flavus*. Overall, this study elucidates the interplay between ecological pressures, antifungal resistance, and genetic differentiation, and invites alternative methods to control aflatoxin contamination in foods without the use of agricultural fungicides.

## Introduction

*Aspergillus flavus* is a ubiquitous, filamentous mold and a significant threat to humans through both food contamination and human infection (1). It is a notorious producer of highly potent, polyketide-derived hepatocarcinogenic aflatoxins that contaminate staple crops including maize and groundnuts (2). Chronic exposure to these aflatoxins is associated with liver cancer, immunosuppression, and other health problems (3). The contamination of agricultural products threatens food safety and economic wellbeing, especially in sub-Saharan Africa, where up to 40% of agricultural production was lost in some regions due to aflatoxin contamination, corresponding to US $450 million (4). In addition, *A. flavus* is the second leading cause of invasive aspergillosis (IA), an often-fatal fungal infection in immunocompromised individuals (5). Environmental conditions such as high temperature and humidity facilitate growth of *A. flavus* and consequently, crop contamination (6). Thus, it is imperative to investigate *A. flavus* populations in regions where the species is particularly prevalent, such as parts of Africa, Asia, and the Middle East, to elucidate important characteristics like genetic structure and antifungal resistance (5). As *A. flavus* is a saprotrophic organism that can produce airborne conidia, it occupies a wide range of ecological niches that provide unique environmental pressures, potentially influencing variations in antifungal susceptibility and genetic diversity (7).

Triazoles are vital to managing *A. flavus*, mitigating infection of human hosts and contamination of crops. Triazoles inhibit fungal growth by targeting the key enzyme lanosterol 14-ɑ demethylase and hindering biosynthesis of ergosterol, a fundamental component of the fungal cell membrane, and increasing intermediate compounds that perturb cell morphology (8). Until recently, *A. flavus* was predominantly susceptible to triazoles, however, emerging studies are challenging this notion. A surveillance study in 2019 reported that 14% of clinical *A. flavus* isolates in South Korea were resistant to voriconazole - a striking increase from the previously determined voriconazole-resistance rate of 0% to 4.9%. It was also found that triazole-resistant strains isolated from patients originated in the environment (9). In *A. flavus*, triazole resistance is primarily associated with mutations in the *cyp51A* gene, which codes for lanosterol demethylase, and overexpression of efflux pumps. However, compared to that in a closely-related species, *Aspergillus fumigatus*, the mechanism of triazole resistance in *A. flavus* has not been widely studied (5). The pervasive application of agricultural triazole fungicides that are structurally similar to medical triazoles favored for the treatment of IA increases the likelihood of cross-resistance, especially as the drugs all share the same target enzyme (10). Nonetheless, limited data are available on antifungal resistance in *A. flavus*, especially in African countries.

Several studies have demonstrated high genetic and genomic diversity in *A. flavus.* For example, one study comparing two isolates of different mating types, with varying stress tolerance, and different aflatoxigenicities, identified more than 100 unique genes and a 310 kilobases (Kb) insertion that contributed to pathogenicity and stress tolerance (11). To explore these findings, Gangurde et al. (12) performed a pangenome analysis of 346 *A. flavus* isolates which supported the accumulation of high genomic diversity during agricultural domestication and emphasized a large accessory genome. Populations also encompass distinct vegetative compatibility (VC) groups from which isolates of the same VC group may anastomose and form stable heterokaryons, enabling transfer of genetic material, contributing to diversity and a complex population structure (13). Nonetheless, reports comparing ecological niche populations are sparse. Recently, Hatmaker et al. (14) analyzed clinical and environmental *A. flavus* populations and found genomic clustering of clinical isolates. However, differences in ecological niche clustering patterns have been reported among studies and such findings suggest that local environmental pressures may influence genetic structure. While the specific mechanisms for such differences are not known, factors such as different selection pressures and the degrees of gene flow and sexual recombination among niches, could contribute to differences in the observed population structures (15, 16).

Despite the widespread impacts of *A. flavus*, there is limited data available regarding the Cameroonian isolates. Current studies focus on a narrow range of ecological sources, as in the study carried out by Nyebe et al. (17), which investigated isolates collected from the rhizosphere or grain of maize, but found no clear correlation between genetic variations and agroecological zone or sample source. Ecological populations from other niches have not been analyzed. Consequently, the extent of genetic variation, antifungal resistance, and niche-specific clustering in Cameroon is poorly understood. As Cameroon’s climate and agricultural practices favor *A. flavus* growth and aflatoxin contamination, comprehensive studies across multiple niches are critical in this species in Cameroon (5, 18).

The objectives of this study were to assess genetic variation and antifungal susceptibility in *A. flavus* isolates obtained from diverse ecological niches in Cameroon, including from groundnut, corn, soil, air, and humans. Using short tandem repeat (STR) markers and triazole susceptibility testing, this study evaluates whether there are niche-specific patterns in genetic structure and triazole resistance. For triazole susceptibility, both clinical and agricultural triazoles were tested. We compared susceptibility profiles across various ecological niche populations and investigated whether different environmental conditions select for distinct genotypes. We hypothesized that triazole resistance will likely be found in Cameroonian *A. flavus* isolates, especially to agricultural triazoles, due to their extensive use to manage the large volume of aflatoxin contamination (19). Exposure to agricultural fungicides selects for strains with mutations that confer resistance, supporting their spread (20). Furthermore, we hypothesized that the minimum inhibitory concentration (MIC) values will correlate with ecological niche, in particular when contrasting agricultural and clinical *A. flavus* isolates. Isolates collected from corn and groundnut were expected to be less susceptible to both fungicides and clinical antifungals because of their high exposure to the fungicides and cross-resistance driven by the structural similarity between the two types of triazoles (10). We also hypothesized that strains belonging to the same niche would genetically cluster because of selective pressures such as competition, nutrient abundance, and moisture levels. In a few studies based in Africa, genetic diversity varied across environmentally distinct regions, and ecological conditions shaped genetic diversity through divergent selective pressures (13, 21). Through the integration of ecological, genetic, and phenotypic data, this research seeks to improve our understanding of *A. flavus* and inform strategies to ameliorate its impact on agriculture and human health.

## Materials and Methods

### Sampling area

The study area is Yaoundé, the political capital and one of the most populated cities in Cameroon with more than 4.5 million inhabitants. Yaoundé city is located at about 250 km east of the Atlantic Ocean and within latitudes 3°50′ and 3°55′ N, and longitudes 11°27′ and 11°35′ E. Its surrounding area is comprised mainly of secondary forests, which are continuously degraded due to subsistence farming. The climate is characterized by annual precipitation of ∼1,600 mm, average temperature of 24° with a long dry season between November and March (22). Sputum and air sampling were performed at the Jamot Hospital, Yaoundé, one of the biggest hospitals in the country that also serves as the main hospital for the management of pulmonary infection in Cameroon. Grains (corn and groundnut) were purchased from two markets and soil samples were collected in three gardens around Yaoundé. The two markets and three gardens were each located at least 1km from each other.

### Sample Collection and Strain Isolation

A total of 560 samples were collected from around Yaoundé, Cameroon, including 100 corn samples, 100 groundnut samples, 100 air samples, 160 soil samples, and 100 clinical samples. These samples were subjected to isolation of *A. flavus* following protocols described previously (23–30). The pure fungal cultures were tentatively identified to *A. flavus* based on their colony morphology and microscopic features. A total of 56 *A. flavus* isolates were obtained, including 23 from corn, 21 from groundnut, 8 from air, and two each from soil samples and human sputum (Table A1). Each isolate was cultured on Sabouraud dextrose agar and incubated at 37℃ for three to six days, until sufficient mycelial growth. The actively growing cultures were then used as inoculants for DNA extraction to be used for STR genotyping and triazole susceptibility testing, following protocols described below.

### DNA extraction and STR Genotyping

DNA extraction was performed according to the phenol-chloroform method described by Xu et al. (31) with minor modifications. Briefly, each isolate was inoculated in Sabouraud dextrose broth at 35℃ for 48 hours. Fungal mycelia were collected from culture, and cell walls were broken down using liquid nitrogen and a pestle. Next, the cells were suspended in a protoplasting buffer, before being centrifuged to collect protoplasts. Lastly, lysing buffer containing Proteinase K was added to each sample, followed by chloroform and isoamyl alcohol, ammonium acetate, isopropyl alcohol, ethanol, and Tris-EDTA buffer, with several mixing steps and pouring of the supernatant in between.

Six STR markers specific to *A. flavus* were analyzed following methods detailed by Halvaeezadeh et al. (32). PCR amplification of these loci utilized fluorescently labeled primers, with the extracted DNA as template (Table A2). The forward primers contained one of the three fluorophores FAM, HEX, or ATTO550 at the 5’ end (Table A3). The primers described by Halvaeezadeh et al. (32) were validated through a manual search of the reference *A. flavus* genome provided by NCBI (RefSeq assembly GCF_009017415.1). The primers for marker *AFPM3* were validated using the NCBI sequence for the locus AY510455 (GenBank). Notably, marker *AFPM4* contained the reverse complement of the dinucleotide repeat ‘CA’, which was listed in the paper as ‘TG’. Because a pilot analysis of amplified fragments indicated that FAM and HEX were not suitable with the primers *AFLA1* and *AFLA7* in a multiplex, HEX was used in singleton reactions to amplify the remaining loci with missing data from the multiplex, facilitating fluorescence detection and fragment analysis (Table A3). The PCR program involved initial denaturation at 94℃ for 15 minutes, followed by 35 cycles of 94℃ for 30 seconds, 54℃ for 30 seconds, and 72℃ for 30 seconds, and a final extension at 72℃ for 10 minutes (32). Amplicons were sent to the Mobix lab at the McMaster Genomics Facility for amplified fragment length polymorphism analysis, from which repeat numbers were derived to infer multilocus genotypes.

### Antifungal Susceptibility Testing

Each isolate was tested for their susceptibilities to two agricultural triazole fungicides, difenoconazole and tebuconazole, and two clinical triazole antifungals, itraconazole and voriconazole. The minimum inhibitory concentration (MIC) was determined for each drug using the 96-well broth microdilution-based protocol outlined by Fan et al. (33). In short, stock solutions of the four antifungals at 3,200 µg/mL were prepared using dimethyl sulfoxide as solvent, and then serially diluted to various working concentrations, before adding RPMI-1640 medium buffered with MOPS. The tested concentrations ranged from 0.015625 µg/mL to 2 µg/mL for the clinical antifungals and 0.03125 µg/mL to 4 µg/mL for the agricultural fungicides, and 0 µg/mL as a control solution. Strains were cultured on Sabouraud dextrose agar plates at 37℃ for 48 hours, including strains of *Candida parapsilosis* (ATCC 90018) and *Candida krusei* (ATCC 14243) for quality controls. After sufficient growth and sporulation, *A. flavus* conidia were harvested and their concentrations were adjusted with a spectrophotometer to an optical density at approximately 0.1 at 530 nm. Strains were then mixed in a ratio of 1:50 with 2xRPMI-1640 and 100 µL of the solution were added to each well containing 100 µL of the corresponding drug. Each strain was incubated in the presence of drug for 48 hours at 35℃. The lowest concentration of each drug at which no visible growth was observed was recorded as the MIC value for the strain. Using the MIC value, each strain was classified as susceptible or resistant based on the European Committee on Antimicrobial Susceptibility Testing (EUCAST) breakpoints and experimental evidence from Jørgensen et al. (34). For voriconazole, isolates with an MIC of ≤ 2 µg/mL were susceptible and those with an MIC of > 2 µg/mL were resistant. For itraconazole, isolates with an MIC value of ≤ 1µg/mL were categorized as susceptible and those with an MIC value of > 2 µg/mL were deemed resistant. Isolates with an MIC value of ≥ 4 µg/mL under difenoconazole or tebuconazole were resistant. Resistant isolates were confirmed through re-testing following the same protocol described above. Moreover, optical density (OD) at 530 nm was used to quantify fungal growth in response to the drugs.

### Computational and Statistical Analysis

OSIRIS was used to score STR genotypes for amplified fragment length polymorphism analysis (35). Specifically, base pair peaks from OSIRIS for each marker were recorded and converted to repeat lengths for downstream analysis. An Analysis of Molecular Variance (AMOVA) test distinguished genetic variation between and within ecological niches using GenAlEx 6.5 (36). Bruvo’s genetic distance, which is specific for STR data, was calculated among strains to perform a Principal Coordinates Analysis (PCoA) using the *poppr* R package (37) and neighbor-joining (NJ) phylogenetic tree, to visualize clustering between strains and ecological niches (38).

Using the MIC data, a Kruskal-Wallis test was performed to compare MIC values between ecological niche samples. To identify which ecological niche populations were significantly different from each other, Wilcoxon rank sum tests with continuity correction and Benjamini-Hochberg adjustments to the p-value, to mitigate false positives, were conducted for pairwise comparisons. OD data was analyzed using a Spearman rank-order correlation test to investigate cross-resistance between triazoles in this population.

Multilocus linkage disequilibrium analyses were performed in Multilocus V1.3. Phylogenetic compatibility and index of association were calculated using formulas as described in the Multilocus V1.3 manual (39).

For all statistical analyses, significance was defined by a p-value of ≤ 0.05. Analyses performed in R used succinct values for MIC results, such that MIC values of “> 2 µg/mL” and “> 4 µg/mL” were treated as “2 µg/mL” and “4 µg/mL,” respectively. This was done to avoid characters that can’t be recognized by the software.

## Results

### Antifungal Susceptibility and Cross-Susceptibility Profiles

Among the 56 *A. flavus* isolates, MIC tests using clinical and agricultural triazoles identified strains resistant to itraconazole, difenoconazole, and/or tebuconazole (Fig. 1). One isolate was resistant to the medical triazole itraconazole (MIC > 2 µg/mL), while 17 (30.3%) isolates were resistant to either difenoconazole or tebuconazole (MIC ≥ 4 µg/mL). Overall, MIC values among isolates in our sample ranged from 0.12 µg/mL to 1 µg/mL for voriconazole, 0.25 µg/mL to > 2 µg/mL for itraconazole, 0.12 µg/mL to 4 µg/mL for difenoconazole, and 0.12 µg/mL to > 4 µg/mL for tebuconazole. Majority of strains were found to be susceptible to all four drugs (39/56; 69.6%). Cross-resistance was observed between agricultural fungicides (11 resistant strains; 19.6%).

**Fig 1.**
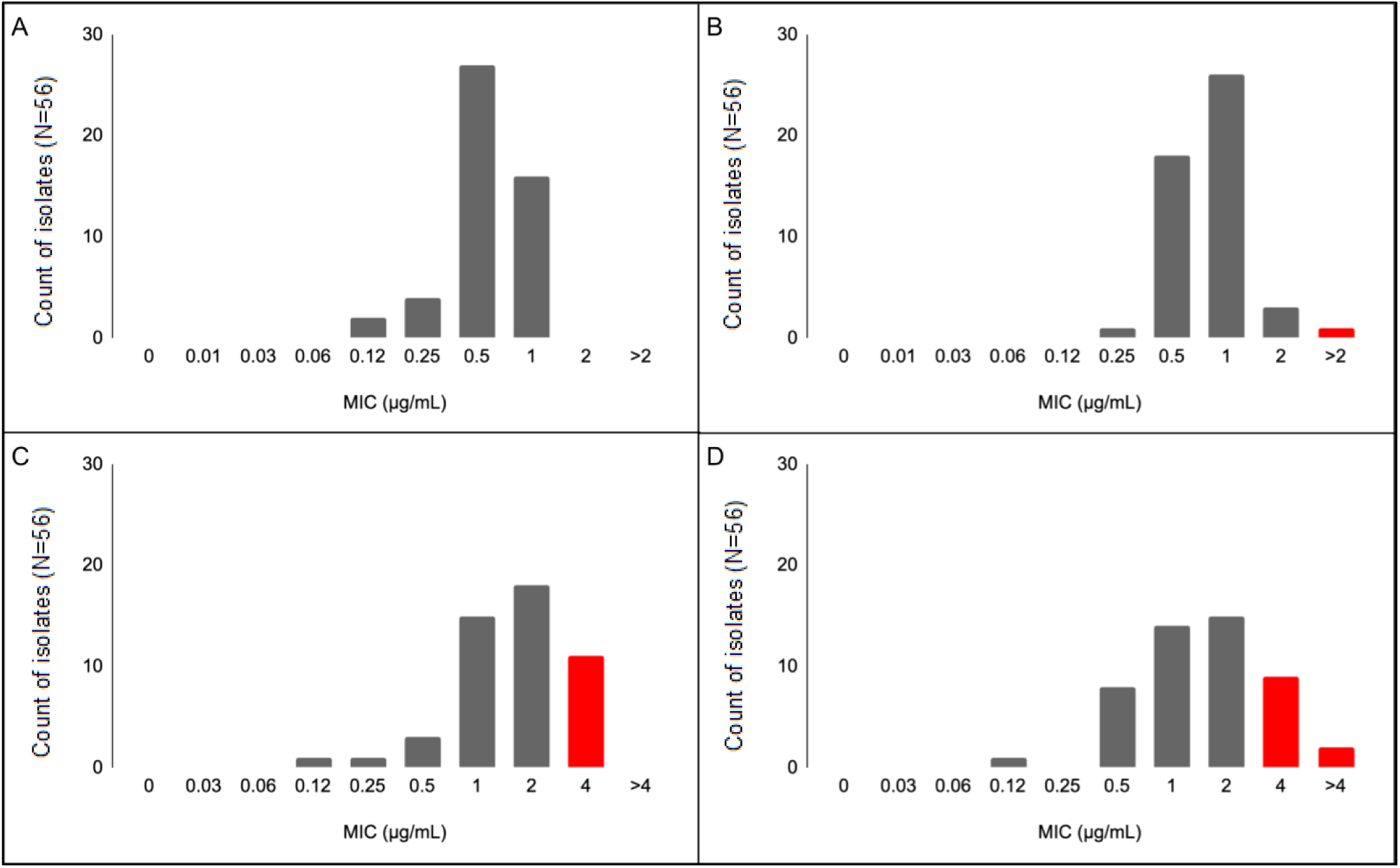
Triazole MIC values among Cameroonian strains of *A. flavus*. One or more strains were resistant under three of the four antifungals tested. MIC testing was performed for 56 isolates against voriconazole, itraconazole (0 to 2 µg/mL), difenoconazole, and tebuconazole (0 to 4 µg/mL). Resistance was classified using EUCAST breakpoints and experimental evidence from Jørgensen et al. (34). Red bars signify resistant isolates. A) No strains were resistant to voriconazole. MIC values ranged from 0.12 to 1 µg/mL among the 56 isolates. B) One isolate was resistant to itraconazole, with an observed fungal growth at 2 µg/mL. MIC values ranged from 0.25 to > 2 µg/mL among the 56 isolates. C) Eleven strains were resistant to difenoconazole, suggested by their growth at 2 µg/mL. MIC values ranged from 0.12 to 4 µg/mL among the 56 isolates. D) Eleven isolates were resistant to tebuconazole, as growth was observed at 2 µg/mL for 9 isolates and at 4 µg/mL for 2 isolates. MIC values ranged from 0.12 to > 4 µg/mL of tebuconazole among the 56 isolates.

To evaluate the impact of ecological niche on antifungal susceptibility, MIC values were compared among isolates from air (n = 8), corn (n = 23), and groundnut (n = 21) environments where sample sizes are relatively large (Fig. 2). Isolates from agricultural crop niches (corn and groundnut) consistently demonstrated elevated MIC values compared to air isolates across both clinical antifungals and agricultural fungicides. This discrepancy was particularly pronounced for agricultural fungicides. For difenoconazole, air isolates exhibited an overall lower MIC profile (median ∼ 1.0 ug/ml) whereas corn and groundnut populations frequently reached the maximum threshold of 4.0 ug/ml. Similarly, tebuconazole MICs were visibly skewed toward higher values in crop-derived isolates, with the corn niche displaying the highest median value (∼ 2.0 ug/ml). Overall, corn and groundnut populations exhibited the widest amounts of variation compared to the air population (0.5 – 4 ug/ml versus 0.5 – 1.25 ug/ml).

**Fig 2.**
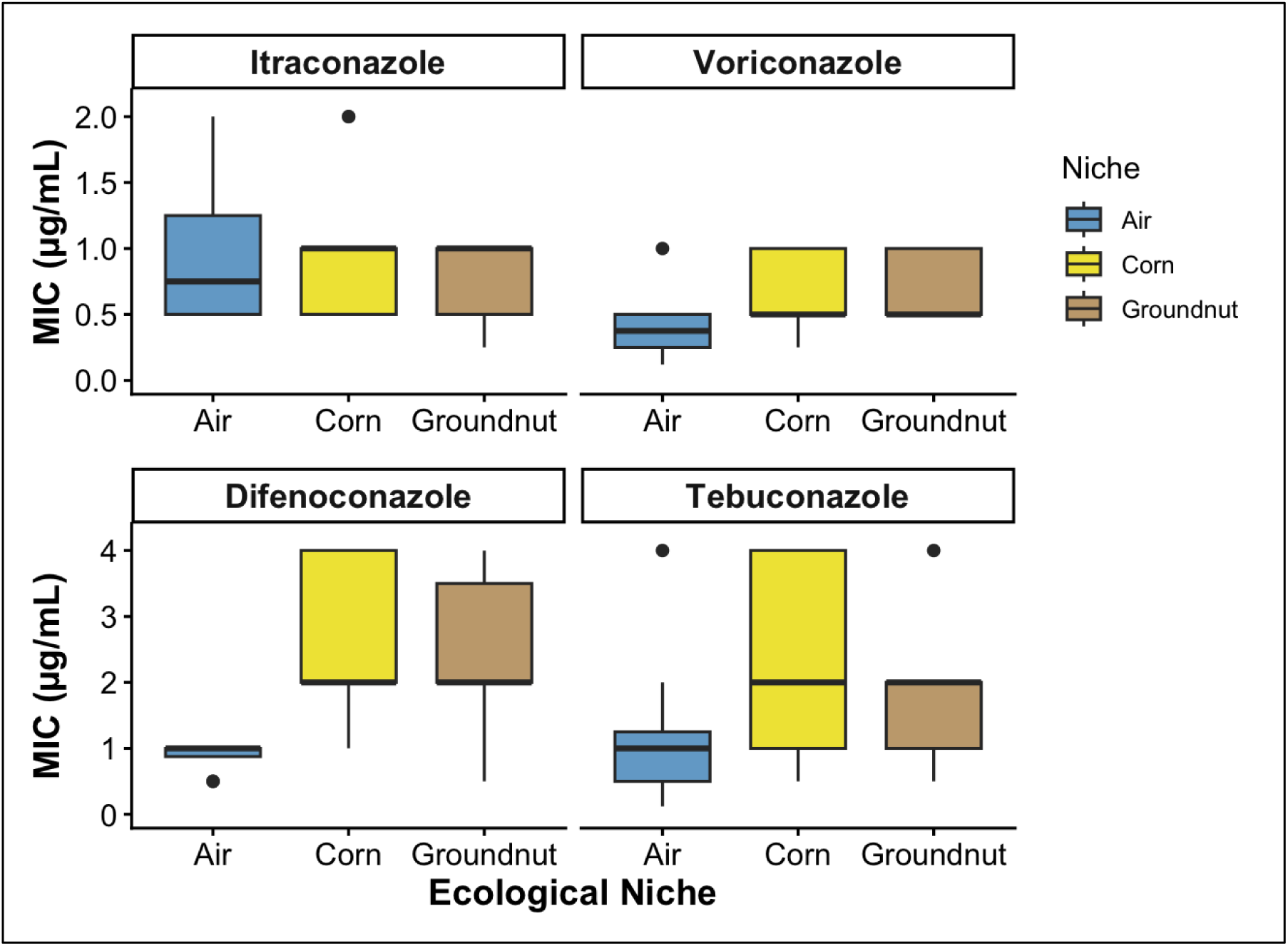
Box plot illustrating the distribution of MIC values across strains from air, corn, and groundnut. MIC values of > 2 µg/mL for clinical antifungals and values of > 4 µg/mL for agricultural fungicides were treated as 2 µg/mL and 4 µg/mL, respectively. MIC testing was performed for 21 groundnut strains, 23 corn strains, 8 air strains, 2 soil isolates, and 2 human isolates. Due to the sample size constraints, populations from soil and humans were excluded from the box plots. Populations from crops show overall increased MIC values compared to air isolates, particularly for voriconazole, difenoconazole, and tebuconazole.

To determine the statistical significance of differences in MIC values between the three niches, a Kruskal-Wallis rank sum test was performed for each drug. For voriconazole, niche-based differences in MIC values were statistically significant (χ^2^=7.81, df=2, p=0.02). Pairwise comparisons using Wilcoxon rank sum test with continuity correction found a significant difference with Benjamini-Hochberg adjusted p-values between corn and air (p=0.04) and groundnut and air (p=0.03). Similarly, analysis of difenoconazole and tebuconazole showed that differences in MIC values were statistically significant (χ^2^=14.22, df=2, p=0.0008 and χ^2^=6.62, df=2, p=0.04, respectively). Under difenoconazole, there was a significant difference between populations from corn and air (p=0.001) and groundnut and air (p=0.002). However, pairwise comparisons of MIC values of tebuconazole found no significant differences between those two compared groups (p > 0.05). Investigation of MIC values for itraconazole revealed that differences in MIC values among the three ecological niches were not statistically significant (χ^2^=1.34, df=2, p=0.51).

As expected, OD data showed that fungal growth decreased overall as antifungal drug concentration increased (Fig. A1). The interquartile range of fungal growth was reduced below the threshold of 0.1 in response to the highest concentration tested of difenoconazole and voriconazole, but not itraconazole or tebuconazole. A Spearman rank-order correlation test between the four triazoles showed strong correlations between all four triazoles (⍴=0.68-0.77; p < 0.001) (Fig. 3). The strongest correlations were observed between the two clinical antifungals and between the two agricultural fungicides, respectively. Moderate correlations were subsequently noted between voriconazole and the agricultural fungicides.

**Fig 3.**
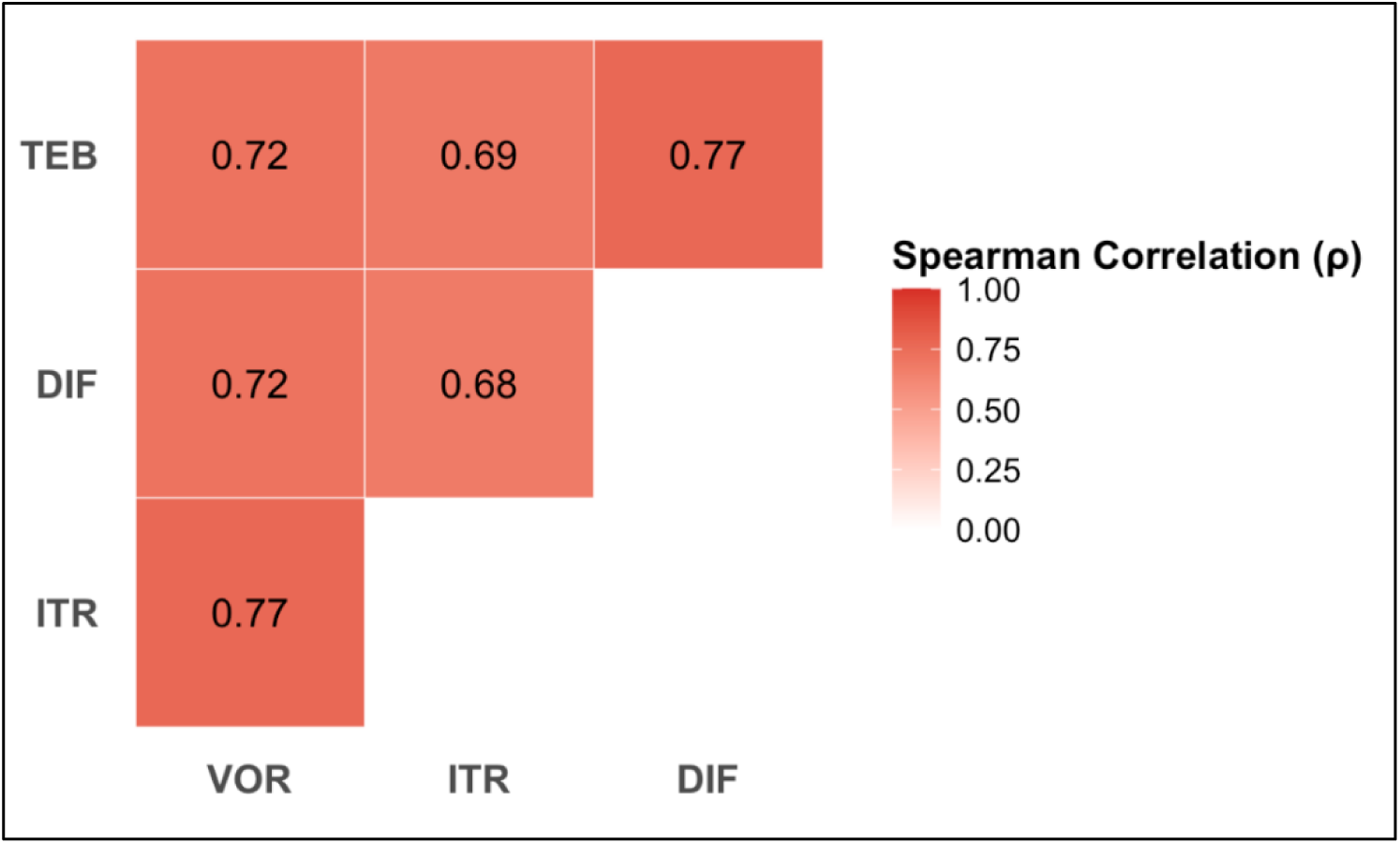
Spearman rank-order correlation test between triazoles shows strong correlations between clinical antifungals and agricultural fungicides. The strongest correlation was observed between voriconazole and itraconazole, and difenoconazole and tebuconazole (⍴=0.77), followed by the fungicides and voriconazole (⍴=0.72), itraconazole and tebuconazole (⍴=0.69), and itraconazole and difenoconazole (⍴=0.68). All correlations were statistically significant (p < 0.001).

### Genetic Differentiation and Population Structure

To determine the partitioning of genetic diversity across different ecological niches, AMOVA was conducted using STR genotypes at six markers analyzed here (Table A4). The analysis revealed that while the vast majority of genetic variation resided within individual ecological niches (82%), a substantial proportion of the total variation was distributed among different ecological niches (18%). The variation between populations from different ecological niches was found to be statistically significant (p<0.001). To visually explore the genetic relationships and differentiation among niches, a PCoA was performed based on Bruvo’s genetic distance, calculated from the six-marker STR genotypes (Fig. 4). The PCoA revealed a distinct clustering pattern corresponding to the ecological niches. Specifically, most isolates from agricultural crop niches (corn and groundnut) clustered tightly along the left side of the first principal coordinate axis (PCoA1), indicating relatively high genetic similarity between isolates in these two ecological populations. In contrast, the air isolates demonstrated a much wider, more dispersed distribution across both axes, reflecting a higher degree of intra-population genetic diversity among the airborne niche isolates.

**Fig 4.**
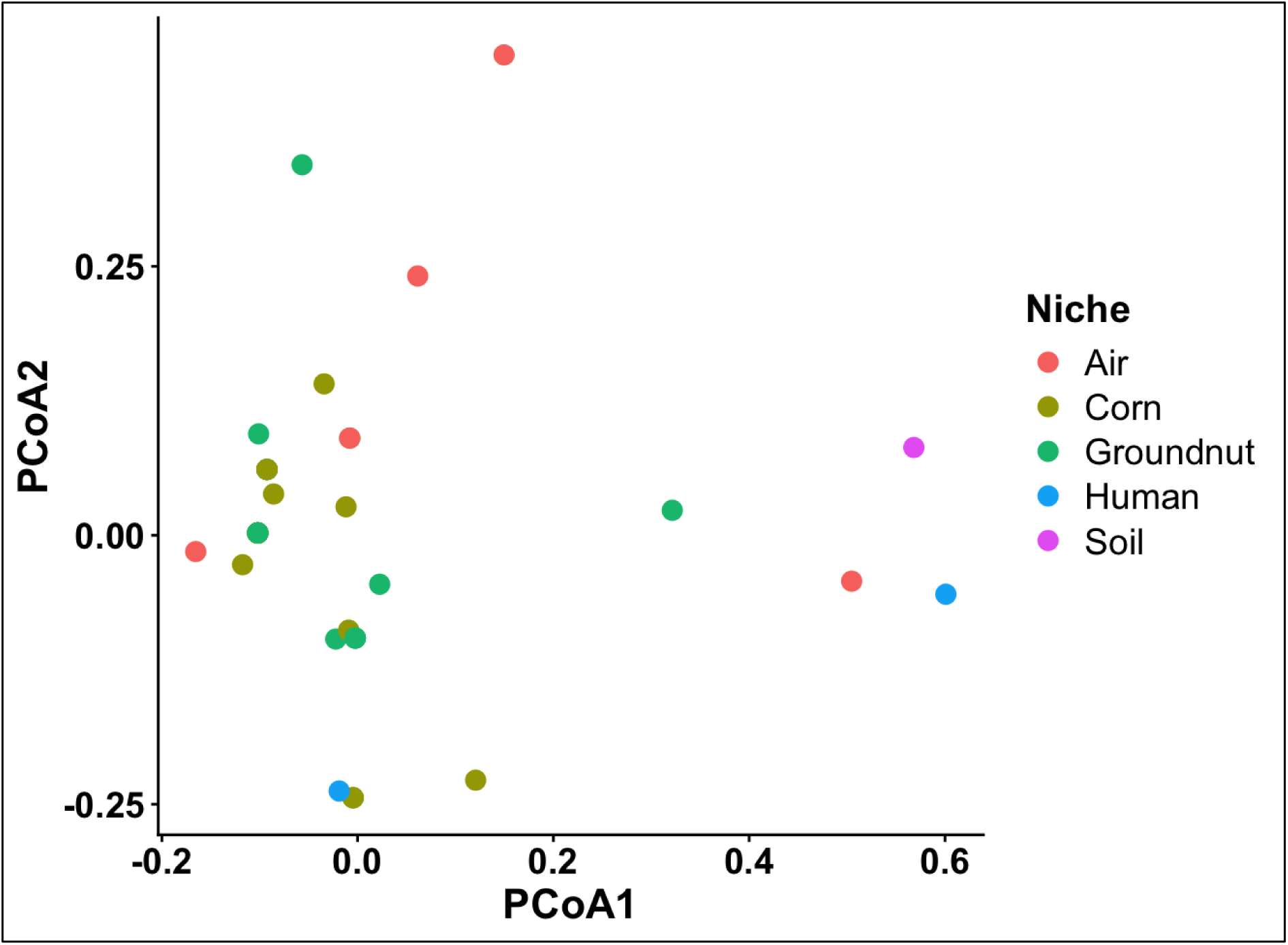
PCoA using Bruvo’s distance visualizes genetic differentiation within and among populations from different ecological niches. STR genotypes were used to calculate Bruvo’s distance between 56 isolates.

To quantify the degree of genetic divergence between specific ecological populations, pairwise phiPTT values were calculated using a Permutational Multivariate Analysis of Variance (PERMANOVA) test across the five ecological niches (Table 1). The analysis revealed varying degrees of population structure, with most pairwise comparisons showing statistically significant genetic differentiation (p < 0.05). The lowest, yet statistically significant, level of differentiation was observed between groundnut and corn populations (phiPTT = 0.048; p=0.024), indicating high genetic similarity between these agricultural populations. In contrast, both crop niches demonstrated highly significant divergence from the airborne population, with groundnut versus air (phiPTT= 0.236; p = 0.001) exhibiting higher differentiation than corn versus air (phiPTT = 0.135; p = 0.004). Due to small sample sizes, the ecological samples from soil and humans were excluded in this analysis.

**Table 1.**
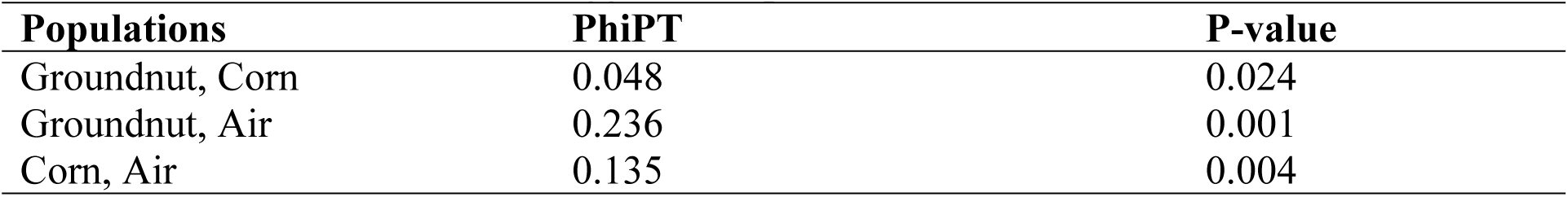
PhiPT from PERMANOVA test, measuring genetic differentiation between populations from different niches. STR genotypes of 21 groundnut strains, 23 corn strains, 8 air strains were analyzed here. A PhiPT value of 0 signifies no differentiation, while a value of 1 suggests complete differentiation.

| Populations | $\phi\text{IPT}$ | P-value |
| --- | --- | --- |
| Groundnut, Corn | 0.048 | 0.024 |
| Groundnut, Air | 0.236 | 0.001 |
| Corn, Air | 0.135 | 0.004 |

### Strain Clustering Analyses

To integrate the genetic and phenotypic datasets into a visual graph, a Neighbor-Joining phylogenetic tree was constructed using Bruvo’s distance and aligned with the corresponding MIC values for clinical antifungals and agricultural fungicides (Fig. 5). The tree demonstrates some ecological niche-based strain clustering, consistent with those observed in the PCoA diagram. Overall, our isolates were grouped into 24 multilocus STR genotypes. Among the 24 multilocus genotypes, 19 were each represented by only one isolate. The remaining five multilocus genotypes were each represented by two to 11 isolates. Four of the five shared multilocus genotypes contained strains from corn and/or groundnuts, while one was shared between air and corn samples, consistent with some level of dispersal between ecological niches. However, almost no two strains with shared multilocus genotypes shared identical triazole susceptibility pattern, consistent with further genetic variation among strains within each of the multilocus genotypes. Indeed, phenotypically, strains with diverse MIC values for each of the four triazoles were overall mixed on the strain relationship tree.

**Fig 5.**
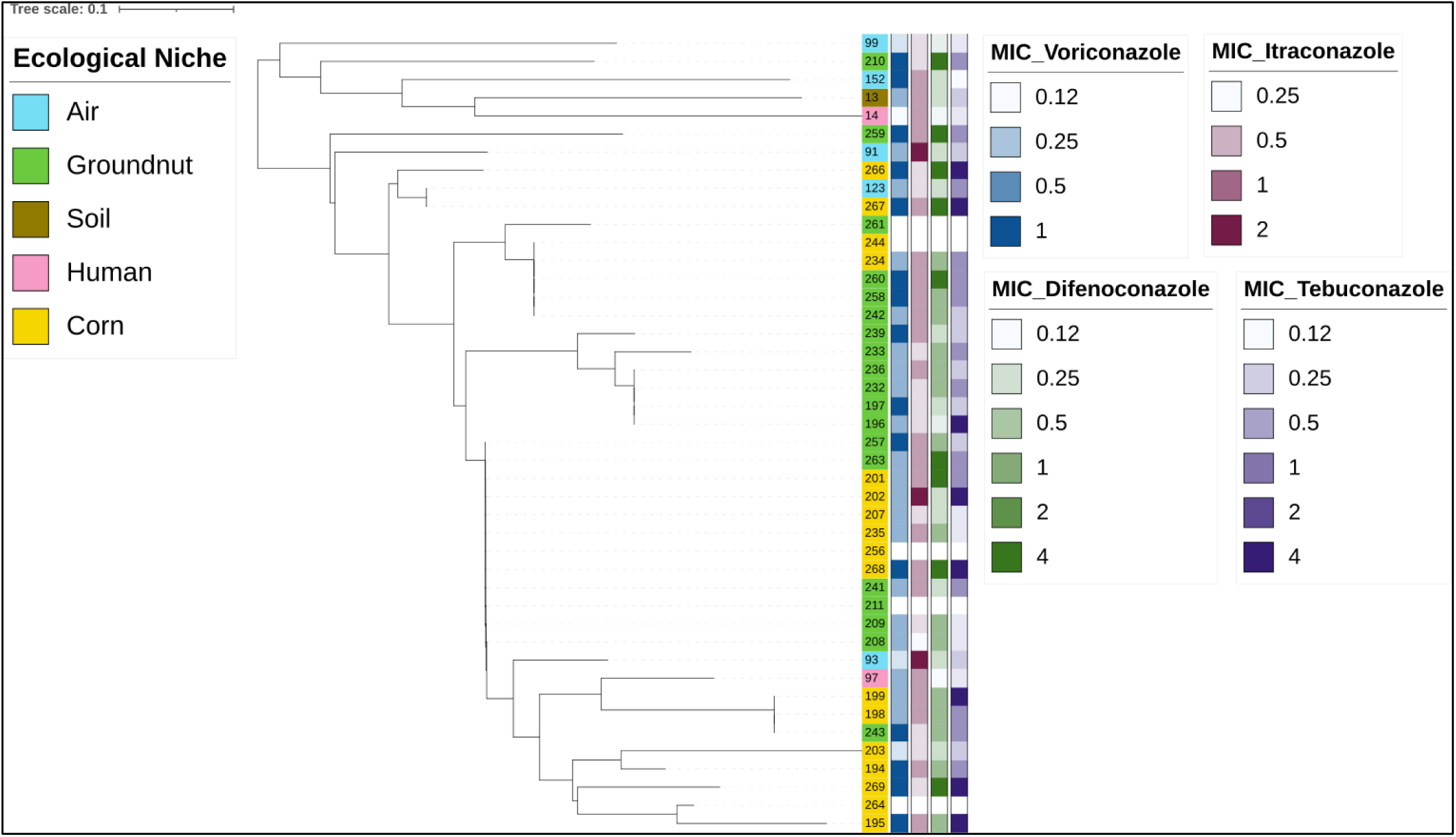
Neighbor-Joining phylogenetic tree based on Bruvo’s distance. Isolate labels are coloured by ecological origin (air, groundnut, soil, human, corn). Adjacent columns depict MICs for voriconazole, itraconazole, difenoconazole, and tebuconazole. Note that MIC values of > 2 µg/mL (clinical antifungals) and > 4 µg/mL (agricultural fungicides) were treated as 2 µg/mL and 4 µg/mL, respectively. STR data was used to calculate Bruvo’s distance and visualized in ITOL for 44 isolates. The remaining 12 isolates were excluded due to deletions in their fragment(s) making them not convertible to Bruvo’s distance (Table A1).

### Evidence for recombination

To investigate whether *A. flavus* populations undergo recombination and if so, whether recombination was random, phylogenetic compatibility and multilocus linkage disequilibrium were evaluated via a permutation test over 999 random replicates (Table A4). Within the total sample, phylogenetic compatibility was 0%, indicating signals of recombination between all pairs of loci (p < 0.01). However, the standardized Index of Association (rBarD) was 0.223 and this value was significantly greater than zero (p < 0.01), rejecting the null hypothesis of random recombination.

## Discussion

In this study, we isolated and analyzed strains of *A. flavus* from five ecological niches Yaounde, Cameroon. Our analyses revealed that while most strains were susceptible to the two agricultural and two medical triazoles that we tested, strains resistant to three of the four triazoles were found, including several strains showing cross-resistance to two antifungals. Overall, samples from corn and groundnut showed higher MIC than those from air. Multilocus genotypic analyses identified high genetic diversity within most ecological niches, with evidence for significant genetic differentiations between air samples and those from corn and groundnut. Interestingly, while several multilocus genotypes were shared between ecological niches, especially between those from corn and groundnut, most strains with shared genotypes had different triazole susceptibility patterns, a result consistent with greater genetic diversity than those revealed by the six STR loci. Below we discuss the relevance of our results to previous studies and their implications to food security and clinical infections.

### Antifungal Susceptibility

In this study, MIC tests were performed to identify patterns of susceptibility to voriconazole, itraconazole, difenoconazole, and tebuconazole in our *A. flavus* ecological populations, and to evaluate the potential impact of ecological niches on antifungal susceptibility. Resistance was observed in response to three of the four triazoles (except voriconazole) for select strains, according to EUCAST breakpoints and experimental evidence from Jørgensen et al. (34) (Fig. 1). While our overall observed triazole resistance rate (30.35%) was high, the rates of resistance to the two medical triazoles were comparable to those reported previously elsewhere, such as those in Iran (20) and Vietnam (40). In both Iran and Vietnam, there were higher rate of resistance to itraconazole than to voriconazole, similar to what we found here. Together, this finding supports EUCAST recommendation in using voriconazole as the primary treatment for invasive aspergillosis by *A. flavus*.

However, it should be noted that EUCAST breakpoints are evolving guidelines. As resistance could be developed due to exposure to agricultural fungicides, and such fungicides are heavily used in Cameroon, resistance to agricultural fungicides could be more common and causing cross-resistance to clinical triazoles (19, 20). In our study, we found a greater number of strains exhibited resistance to agricultural fungicides compared to clinical antifungals (Fig. 1). In general, a higher concentration of difenoconazole and tebuconazole were needed to inhibit *A. flavus* growth compared to clinical antifungals (Fig. 1). Together, this emphasizes the need for regulations on fungicide use in Cameroon and a potential transition to different classes of fungicides or to biocontrol agents, to both control aflatoxin contamination of agricultural products and prevent further development of antifungal resistance.

Aside from MIC determinations, we also quantified fungal growth in response to each of the triazoles. Our analyses identified voriconazole and difenoconazole as the superior clinical antifungal and agricultural fungicide, respectively. Indeed, some strains of *A. flavus* showed vigorous growth at the highest concentration that we tested for itraconazole and tebuconazole (Fig. A1). OD measurements also indicate the potential of cross-resistance between the four triazoles (Fig. 3). Indeed, fungal growth under all four drugs were strongly correlated, with itraconazole and voriconazole showing the strongest correlation, followed by difenoconazole and tebuconazole. Notably, fungal growth in response to voriconazole was also strongly correlated with growth under agricultural fungicides (⍴=0.72), and similarly, growth was strongly correlated between itraconazole and the agricultural fungicides (⍴=0.68-0.69). As fungicide use is pervasive in Cameroon, these findings heighten the possibility of tolerance and resistance to both agricultural and clinical triazoles, especially due to the chemical structural similarity and similar mechanism of action of the two types of triazoles (19, 41). While resistance to medical triazoles in patients due to exposure to agricultural fungicides has yet to be reported in *A. flavus*, given the close genetic relatedness between clinical and crop isolates, our study may represent the first evidence that such a development can happen in this species (5). As fungicides continue to be applied in agricultural practices to reduce aflatoxin contamination, these drugs may become less effective in agricultural and clinical settings, increasing the risk of fatality in individuals with invasive aspergillosis due to the emergence of voriconazole- and itraconazole-resistant *A. flavus* strains.

Through a Kruskal-Wallis rank sum test, it was found that differences in MIC values between the three ecological niches were statistically significant in response to voriconazole, difenoconazole, and tebuconazole (p < 0.05). Specifically, isolates from crops were overall less susceptible to voriconazole and difenoconazole than isolates collected from air (Fig. 2). Further pairwise comparisons revealed several niche-based clustering of isolates in their antifungal susceptibility, specifically between some crop-associated isolates (groundnut, corn) and air in response to voriconazole and difenoconazole (p < 0.05). These findings support the hypothesis that antifungal susceptibility varies across niches, with agricultural isolates displaying less susceptibility due to fungicide use. Because the air population of *A. flavus* represents the airborne portion of other ecological populations such as from the soil, its overall lower MICs than those from groundnut and corn in Yaoundé suggest that at least in the hospital environment where air samples were collected, the air samples seemed to have been minimally impacted by spores from infected groundnuts and corn. Studies in Asia and the Middle East report higher resistance rates in environmental isolates compared to clinical isolates as well, supporting the notion that exposure to fungicides may lead to acquired resistance in *A. flavus.* Specifically, the application of fungicides in agricultural regions selects for strains that have mutations in the *cyp51* gene, increasing the prevalence of resistant strains (20).

### Genetic Variation

Our population genetic analyses of STR genotypes revealed that 18% of the total genetic variation was attributed by ecological niche separations (p < 0.001). In the PCoA using Bruvo’s genetic distance, there was some clustering of corn and groundnut strains (Fig. 4). The most genetically different populations were isolates from groundnut and human samples whereas the most similar populations were between groundnut and corn. This is likely due to the similar ecological pressures faced by crops and crop-associated pathogens, and the vastly different environments of a human host compared to an agricultural product. In fungi, biosynthetic gene clusters have been found to be involved in niche-based clustering as they may be selected for in specific ecological conditions and affect which niches the fungi can colonize (42). Whole genome sequencing could help identify such niche-adaptation associated genes in our samples.

Giving the ability of *A. flavus* to reproduce asexually, the lack of tight associations between genotypes and antifungal susceptibilities was surprising (Fig. 5). This result suggests greater genotypic diversity among our isolates than what we revealed using the six STR markers. In a study in Iran that used nine STR markers, most isolates had different genotypes (5). The high diversity of genotypes is consistent with recombination in natural populations of *A. flavus*. Indeed, multilocus linkage disequilibrium analysis revealed evidence for recombination between all six STR loci in the *A. flavus* population in Cameroon. Evidence of recombination has been observed before for *A. flavus* (43). Together, increasing data support recombination playing an important role in natural populations of *A. flavus*.

### Limitations and Perspectives

For both antifungal susceptibility and genetic variation analyses, niche-based comparisons are limited in their statistical power due to low sample sizes for certain populations. Overall, despite relatively large ecological samples that we processed, only two isolates of *A. flavus* from each of the soil and humans were obtained. For this reason, the soil and human samples were excluded from statistical analyses in niche-based population genetic comparisons. Despite the sample size limitation, our analyses found overall high similarities in both genotypes and triazole susceptibility patterns between *A. flavus* samples from corn and groundnut, likely due to similar management practices after their harvests.

In conclusion, the presence of resistant strains to itraconazole, difenoconazole, and tebuconazole, coupled by the lack of strains resistant to voriconazole support voriconazole as the primary treatment for IA caused by *A. flavus*. However, the identification of significant correlations in fungal growth under various agriculture and clinical triazoles indicates a worrisome trend that could impact future treatment of *A. flavus* infections by triazoles. Our data highlight the need for more stringent controls of application of agricultural fungicides and a One Health approach to effectively prevent and control *A. flavus* infections in both agriculture and the clinics (44).

## Acknowledgments

We thank the staff at Jamot Hospital, Yaoundé, Cameroon for helping with patient and air sample collections. We thank the staff at Molecular Biology Center, Yaoundé, Cameroon for assistance in isolating *A. flavus* from various samples.

## Author contributions

Conceptualization: MK, JX. Investigations: MK, KR, JD, JN, JH, CN, RN, JX. Funding acquisition: JX, MK. Project administration: JX. Supervision: JX, CN, RN. Writing – original draft: MK, KR, and JD. Writing – review & editing: JD, JX.

## Data availability

All data supporting the conclusions of the study are presented in the manuscript.

## Appendix

**Table A1.**
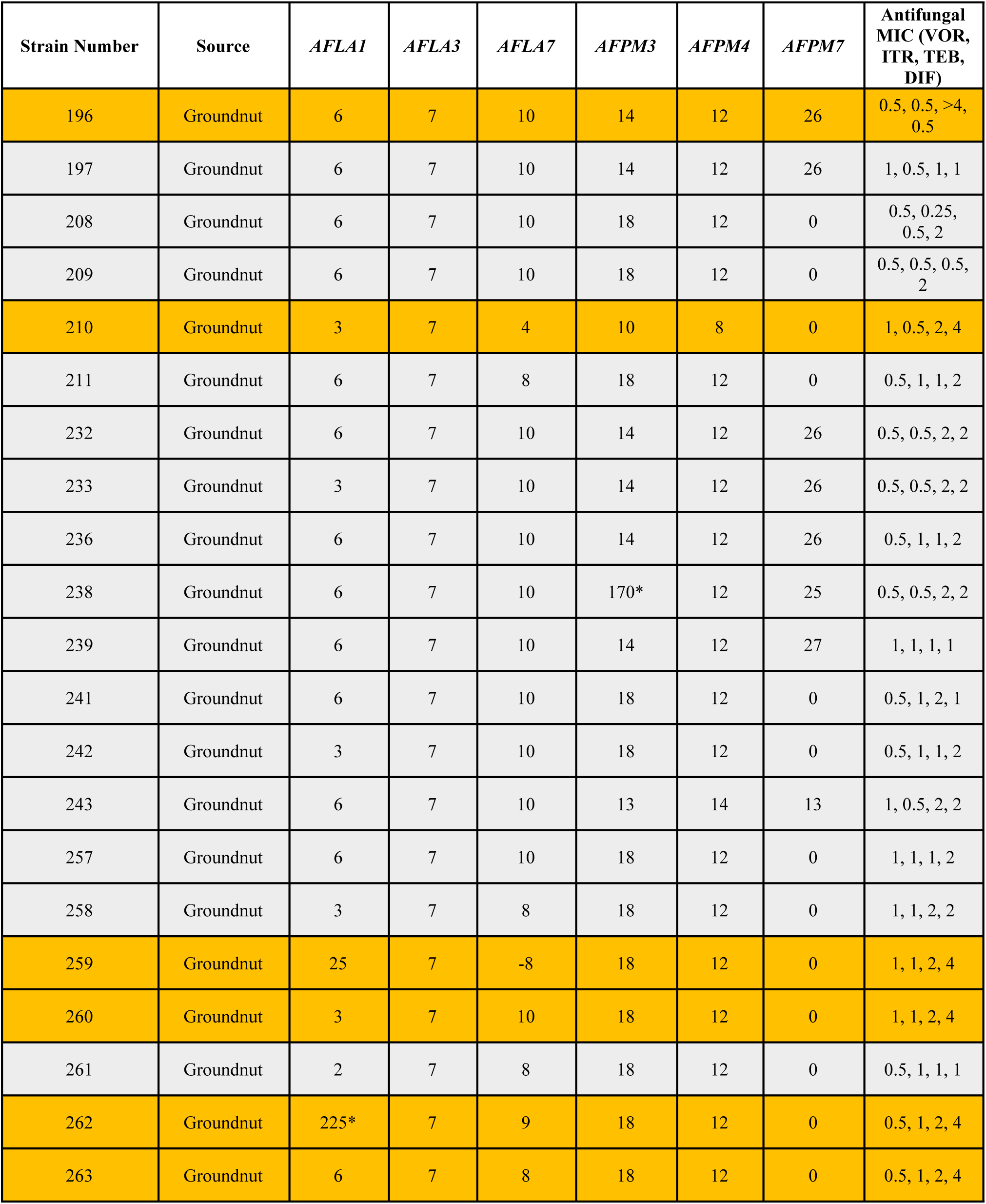
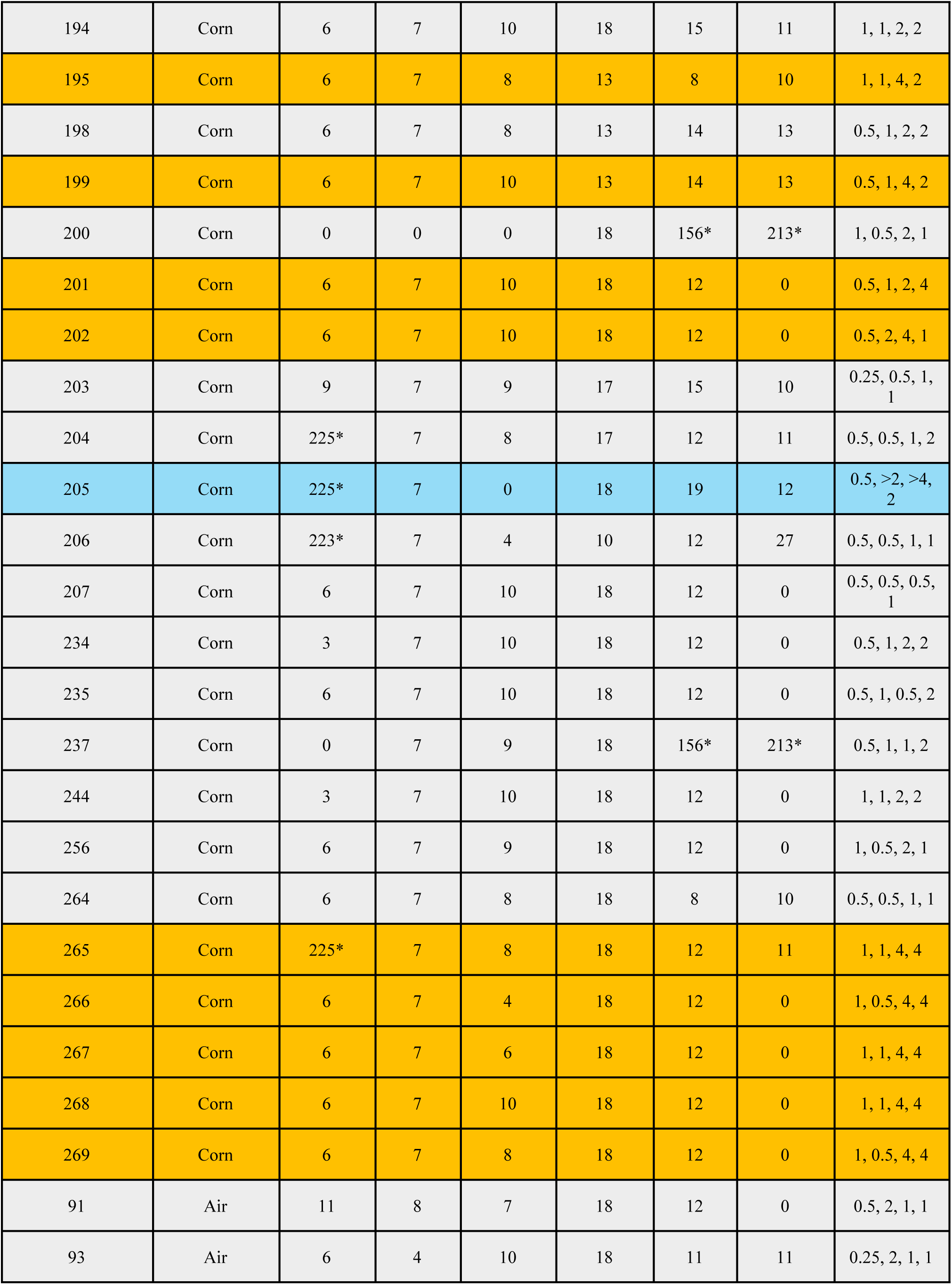
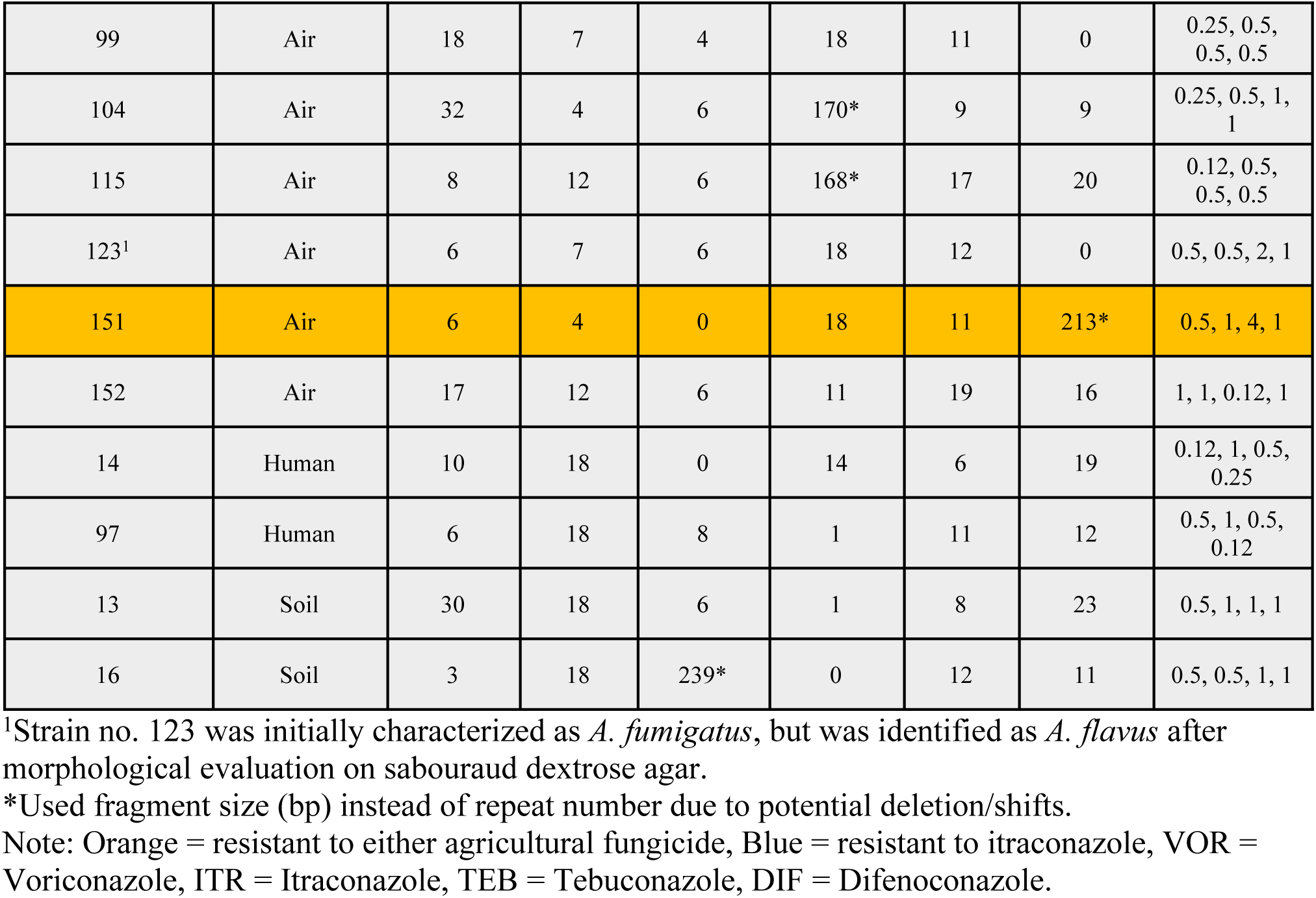
Overview of *Aspergillus flavus (A. flavus)* isolates collected from Yaoundé, Cameroon.

| Strain Number | Source | *AFLA1* | *AFLA3* | *AFLA7* | *AFPM3* | *AFPM4* | *AFPM7* | Antifungal MIC (VOR, ITR, TEB, DIF) |
| --- | --- | --- | --- | --- | --- | --- | --- | --- |
| 196 | Groundnut | 6 | 7 | 10 | 14 | 12 | 26 | 0.5, 0.5, >4, 0.5 |
| 197 | Groundnut | 6 | 7 | 10 | 14 | 12 | 26 | 1, 0.5, 1, 1 |
| 208 | Groundnut | 6 | 7 | 10 | 18 | 12 | 0 | 0.5, 0.25, 0.5, 2 |
| 209 | Groundnut | 6 | 7 | 10 | 18 | 12 | 0 | 0.5, 0.5, 0.5, 2 |
| 210 | Groundnut | 3 | 7 | 4 | 10 | 8 | 0 | 1, 0.5, 2, 4 |
| 211 | Groundnut | 6 | 7 | 8 | 18 | 12 | 0 | 0.5, 1, 1, 2 |
| 232 | Groundnut | 6 | 7 | 10 | 14 | 12 | 26 | 0.5, 0.5, 2, 2 |
| 233 | Groundnut | 3 | 7 | 10 | 14 | 12 | 26 | 0.5, 0.5, 2, 2 |
| 236 | Groundnut | 6 | 7 | 10 | 14 | 12 | 26 | 0.5, 1, 1, 2 |
| 238 | Groundnut | 6 | 7 | 10 | 170* | 12 | 25 | 0.5, 0.5, 2, 2 |
| 239 | Groundnut | 6 | 7 | 10 | 14 | 12 | 27 | 1, 1, 1, 1 |
| 241 | Groundnut | 6 | 7 | 10 | 18 | 12 | 0 | 0.5, 1, 2, 1 |
| 242 | Groundnut | 3 | 7 | 10 | 18 | 12 | 0 | 0.5, 1, 1, 2 |
| 243 | Groundnut | 6 | 7 | 10 | 13 | 14 | 13 | 1, 0.5, 2, 2 |
| 257 | Groundnut | 6 | 7 | 10 | 18 | 12 | 0 | 1, 1, 1, 2 |
| 258 | Groundnut | 3 | 7 | 8 | 18 | 12 | 0 | 1, 1, 2, 2 |
| 259 | Groundnut | 25 | 7 | -8 | 18 | 12 | 0 | 1, 1, 2, 4 |
| 260 | Groundnut | 3 | 7 | 10 | 18 | 12 | 0 | 1, 1, 2, 4 |
| 261 | Groundnut | 2 | 7 | 8 | 18 | 12 | 0 | 0.5, 1, 1, 1 |
| 262 | Groundnut | 225* | 7 | 9 | 18 | 12 | 0 | 0.5, 1, 2, 4 |
| 263 | Groundnut | 6 | 7 | 8 | 18 | 12 | 0 | 0.5, 1, 2, 4 |
| 194 | Corn | 6 | 7 | 10 | 18 | 15 | 11 | 1, 1, 2, 2 |
| 195 | Corn | 6 | 7 | 8 | 13 | 8 | 10 | 1, 1, 4, 2 |
| 198 | Corn | 6 | 7 | 8 | 13 | 14 | 13 | 0.5, 1, 2, 2 |
| 199 | Corn | 6 | 7 | 10 | 13 | 14 | 13 | 0.5, 1, 4, 2 |
| 200 | Corn | 0 | 0 | 0 | 18 | 156* | 213* | 1, 0.5, 2, 1 |
| 201 | Corn | 6 | 7 | 10 | 18 | 12 | 0 | 0.5, 1, 2, 4 |
| 202 | Corn | 6 | 7 | 10 | 18 | 12 | 0 | 0.5, 2, 4, 1 |
| 203 | Corn | 9 | 7 | 9 | 17 | 15 | 10 | 0.25, 0.5, 1,<br>1 |
| 204 | Corn | 225* | 7 | 8 | 17 | 12 | 11 | 0.5, 0.5, 1, 2 |
| 205 | Corn | 225* | 7 | 0 | 18 | 19 | 12 | 0.5, >2, >4,<br>2 |
| 206 | Corn | 223* | 7 | 4 | 10 | 12 | 27 | 0.5, 0.5, 1, 1 |
| 207 | Corn | 6 | 7 | 10 | 18 | 12 | 0 | 0.5, 0.5, 0.5,<br>1 |
| 234 | Corn | 3 | 7 | 10 | 18 | 12 | 0 | 0.5, 1, 2, 2 |
| 235 | Corn | 6 | 7 | 10 | 18 | 12 | 0 | 0.5, 1, 0.5, 2 |
| 237 | Corn | 0 | 7 | 9 | 18 | 156* | 213* | 0.5, 1, 1, 2 |
| 244 | Corn | 3 | 7 | 10 | 18 | 12 | 0 | 1, 1, 2, 2 |
| 256 | Corn | 6 | 7 | 9 | 18 | 12 | 0 | 1, 0.5, 2, 1 |
| 264 | Corn | 6 | 7 | 8 | 18 | 8 | 10 | 0.5, 0.5, 1, 1 |
| 265 | Corn | 225* | 7 | 8 | 18 | 12 | 11 | 1, 1, 4, 4 |
| 266 | Corn | 6 | 7 | 4 | 18 | 12 | 0 | 1, 0.5, 4, 4 |
| 267 | Corn | 6 | 7 | 6 | 18 | 12 | 0 | 1, 1, 4, 4 |
| 268 | Corn | 6 | 7 | 10 | 18 | 12 | 0 | 1, 1, 4, 4 |
| 269 | Corn | 6 | 7 | 8 | 18 | 12 | 0 | 1, 0.5, 4, 4 |
| 91 | Air | 11 | 8 | 7 | 18 | 12 | 0 | 0.5, 2, 1, 1 |
| 93 | Air | 6 | 4 | 10 | 18 | 11 | 11 | 0.25, 2, 1, 1 |
| 99 | Air | 18 | 7 | 4 | 18 | 11 | 0 | 0.25, 0.5, 0.5, 0.5 |
| 104 | Air | 32 | 4 | 6 | 170* | 9 | 9 | 0.25, 0.5, 1, 1 |
| 115 | Air | 8 | 12 | 6 | 168* | 17 | 20 | 0.12, 0.5, 0.5, 0.5 |
| 123<sup>1</sup> | Air | 6 | 7 | 6 | 18 | 12 | 0 | 0.5, 0.5, 2, 1 |
| 151 | Air | 6 | 4 | 0 | 18 | 11 | 213* | 0.5, 1, 4, 1 |
| 152 | Air | 17 | 12 | 6 | 11 | 19 | 16 | 1, 1, 0.12, 1 |
| 14 | Human | 10 | 18 | 0 | 14 | 6 | 19 | 0.12, 1, 0.5, 0.25 |
| 97 | Human | 6 | 18 | 8 | 1 | 11 | 12 | 0.5, 1, 0.5, 0.12 |
| 13 | Soil | 30 | 18 | 6 | 1 | 8 | 23 | 0.5, 1, 1, 1 |
| 16 | Soil | 3 | 18 | 239* | 0 | 12 | 11 | 0.5, 0.5, 1, 1 |

<sup>1</sup>Strain no. 123 was initially characterized as *A. fumigatus*, but was identified as *A. flavus* after morphological evaluation on sabouraud dextrose agar.
\*Used fragment size (bp) instead of repeat number due to potential deletion/shifts.
Note: Orange = resistant to either agricultural fungicide, Blue = resistant to itraconazole, VOR = Voriconazole, ITR = Itraconazole, TEB = Tebuconazole, DIF = Difenconazole.

**Table A2.**
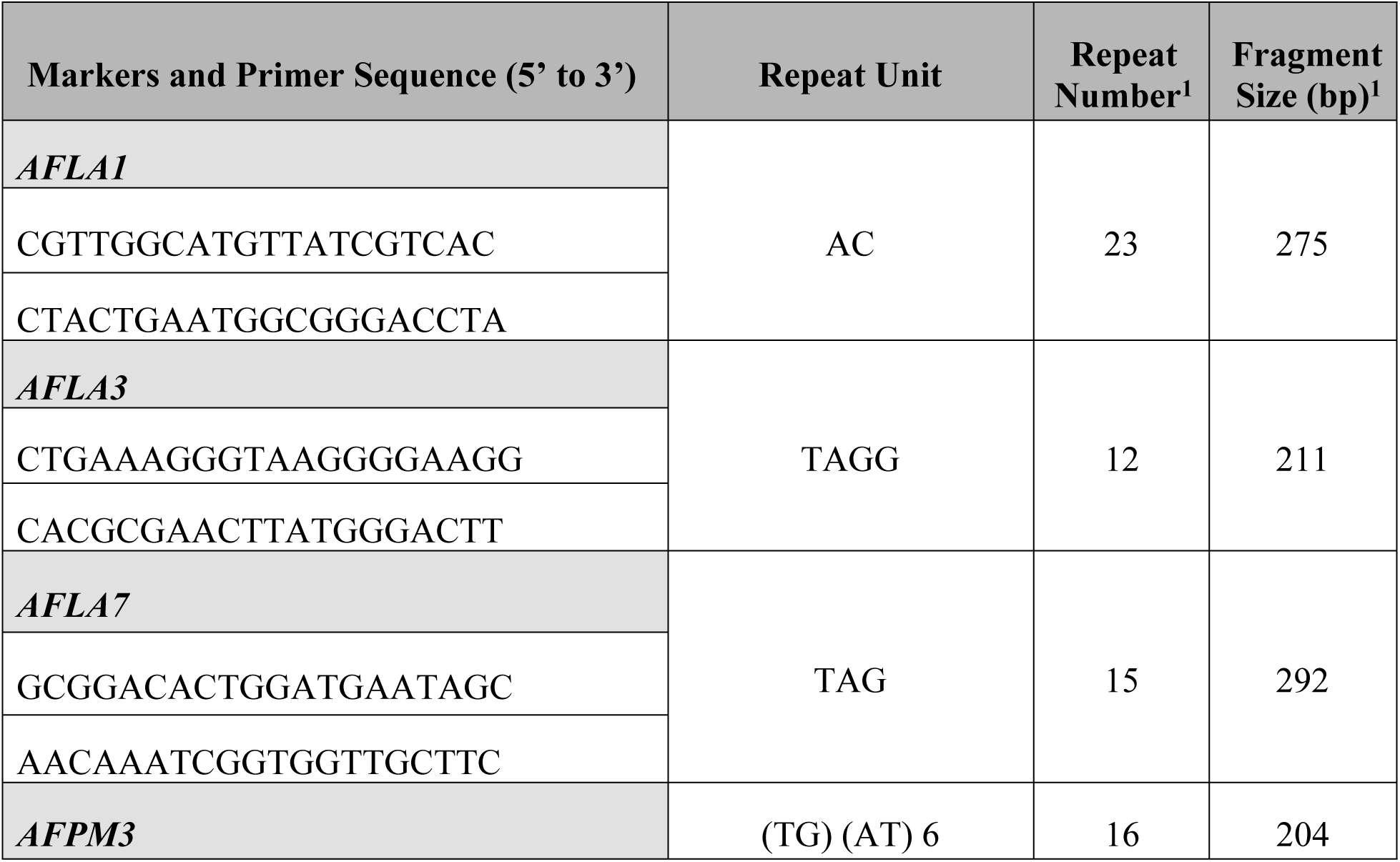
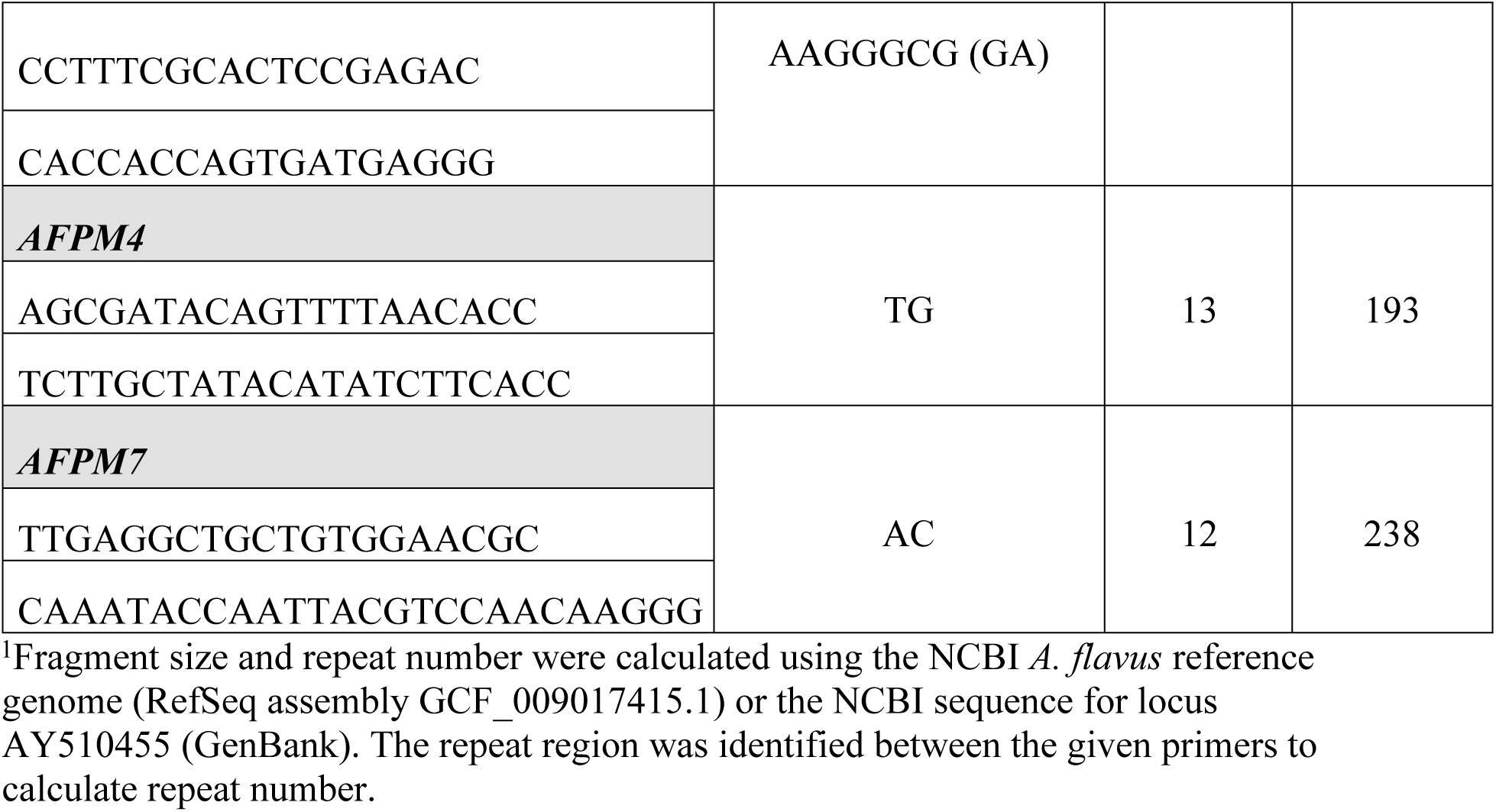
Microsatellite Length Polymorphism Markers and Characteristics (modified from Halvaeezadeh et al., 2025); (32)

**Table A3.**
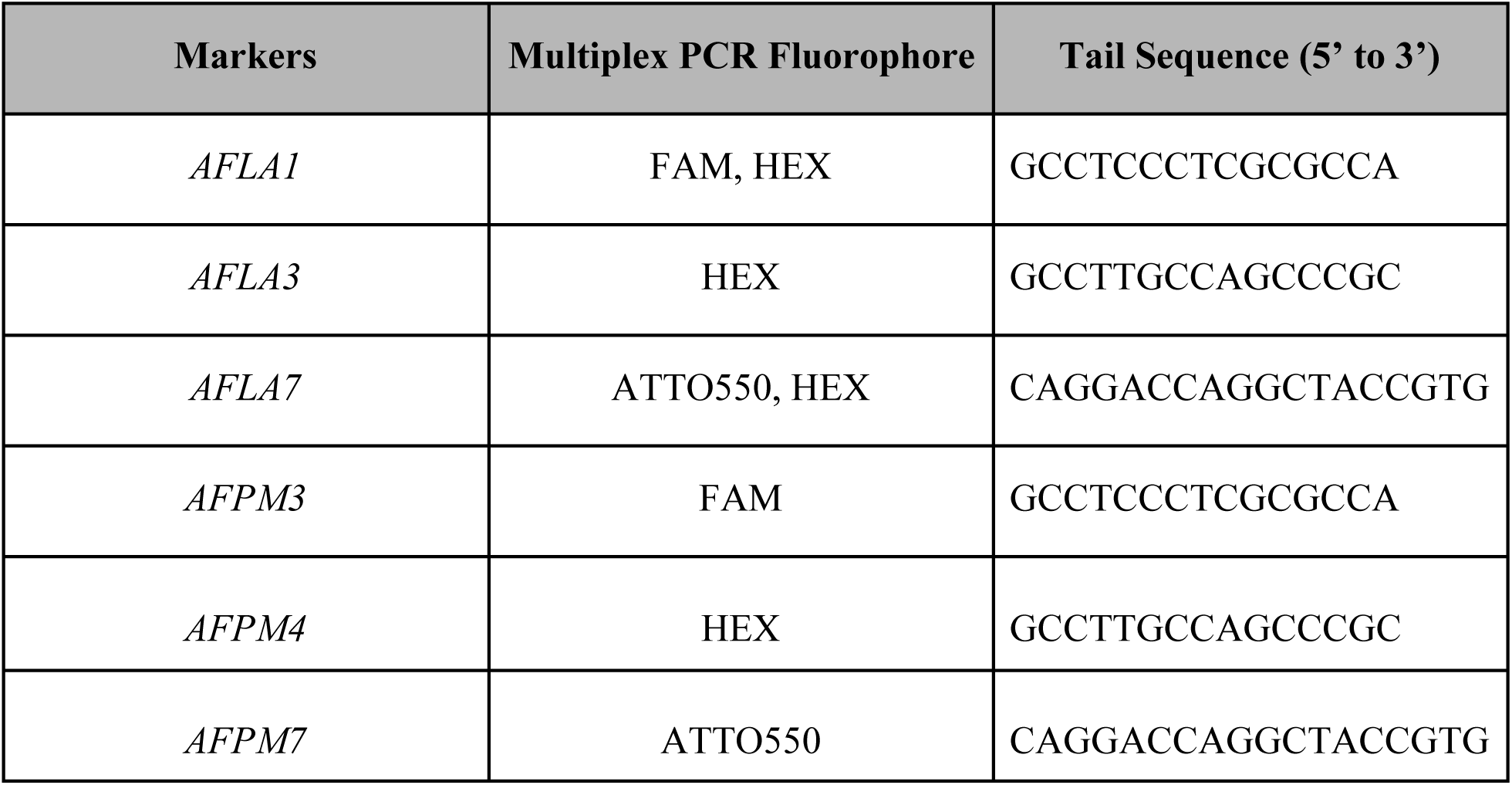
Multiplex PCR Fluorophores and Tail Sequences (modified from Blacket et al., 2012); (43).

**Table A4.**
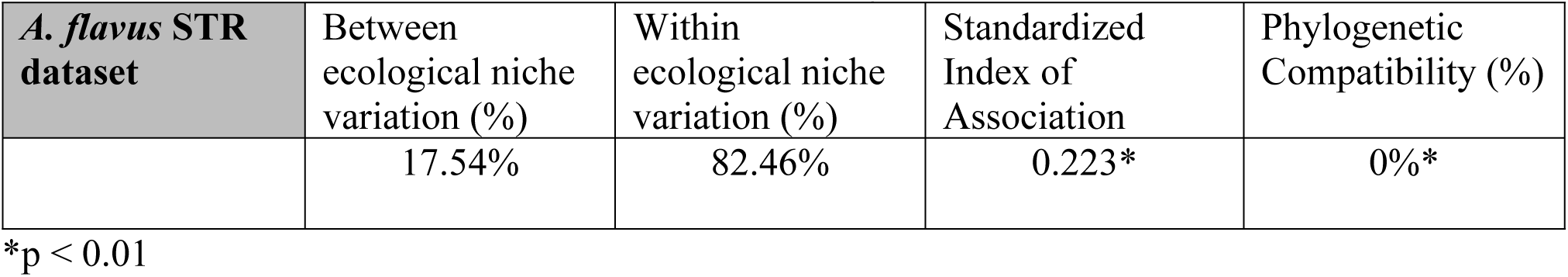
Analysis of molecular variance and linkage disequilibrium analysis among Cameroonian *A. flavus*.

**Figure A1.**
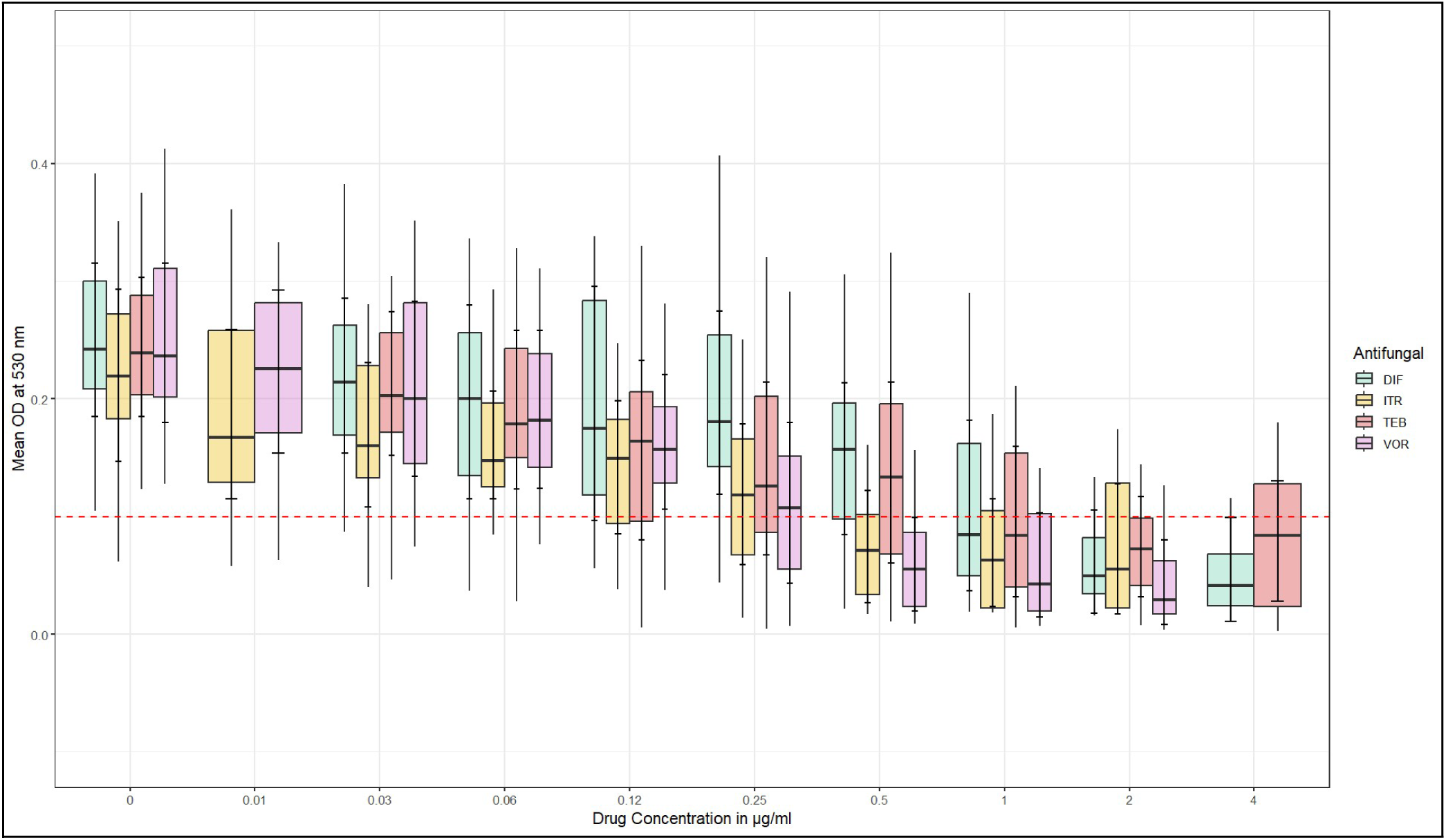
Box plot illustrating fungal growth across different concentrations of difenoconazole, itraconazole, tebuconazole, and voriconazole. Growth was measured using optical density measured at 530 nm. Red line (y=0.1) represents a threshold for growth. The interquartile range of fungal growth was reduced below the threshold in response to the highest concentration tested of difenoconazole and voriconazole, but not itraconazole or tebuconazole.

## Calculation of Repeat Number from Fragment Size

Fragment size was identified for each marker on OSIRIS. Differences between the fragment size found in the isolate of study and the reference sequence were calculated. Using the size difference, the change in repeat number was found, which was used to determine the true repeat number in the isolate of study for each marker.

## M1-FAM (strain 196)

Fragment Size on OSIRIS = 241

M1-FAM corresponds to marker *AFLA1*, which has a fragment size of 275 (Table. 2)

Fragment Size Difference = 275-241 = 34

*AFLA1* contains a dinucleotide repeat, thus 34 less base pairs in the isolate of study

corresponds to 17 fewer repeats (34/2=17) (Table. 2)

No. of Repeats in Reference = 23 (Table. 2)

No. of Repeats in Strain 196 = 23-17 = 6

Thus, isolate 196 contains 6 repeats at the locus *AFLA1*

**Table A5.** Summary of results from STR genotyping and MIC testing. Characteristics of six microsatellite markers and MIC results for clinical and agricultural triazoles distributed across niche. All available data was compiled to determine the number of unique alleles per niche.

| Markers | Fragment Size Range (bp) | Repeat No. Range | No. of Unique Alleles |  |  |  |  |  |
| --- | --- | --- | --- | --- | --- | --- | --- | --- |
|  |  |  | Corn | Groundnut | Air | Soil | Human | Total |
| *AFLA1* | 234 - 294 | 2 - 32 | 3 | 4 | 6 | 2 | 2 | 10 |
| *AFLA3* | 181 - 234 | 4 - 18 | 1 | 1 | 4 | 1 | 1 | 5 |
| *AFLA7* | 224 - 277 | -8 - 10 | 5 | 5 | 4 | 2 | 2 | 8 |
| *AFPM3* | 174 - 208 | 1 - 18 | 4 | 4 | 2 | 1 | 2 | 7 |
| *AFPM4* | 156 - 204 | -5 - 19 | 6 | 3 | 6 | 2 | 2 | 8 |
| *AFPM7* | 212 - 269 | -1 - 27 | 6 | 6 | 6 | 2 | 2 | 11 |
| MIC Range (µg/mL) | Clinical | Voriconazole | 0.25 - 1 | 0.5 - 1 | 0.12 - 1 | 0.5 | 0.12 - 0.5 | - |
|  |  | Itraconazole | 0.5 - > 2 | 0.25 - 1 | 0.5 - 2 | 0.5 - 1 | 1 |  |
|  | Agricultural | Tebuconazole | 0.5 - > 4 | 0.5 - > 4 | 0.12 - 4 | 1 | 0.5 |  |
|  |  | Difenoconazole | 1 - 4 | 0.5 - 4 | 0.5 - 1 | 1 | 0.12 – 0.25 |  |
| Total No. of Isolates |  |  | 23 | 21 | 8 | 2 | 2 | 56 |

## References

1. Xu J. Assessing Global Fungal Threats to Humans. mLife. 2022;1(3): 223–240. 10.1002/mlf2.12036

2. Abdullaziz S, Zhuang Y, Zhang C, Sharif Y, Ibrahim MM, Nassarawa IS, et al. Double-edged sword: *Aspergillus flavus* as a threat to food safety and a resource for biotechnology. Biotechnol. Adv., 2026;89: 108848. 10.1016/j.biotechadv.2026.108848

3. Fagbohun TR, Nji QN, Okechukwu VO, Adelusi OA, Nyathi LA, Awong P, et al. Aflatoxin Exposure in Immunocompromised Patients: Current State and Future Perspectives. Tox. 2015;17(8): 414. 10.3390/toxins17080414

4. Saber H, Chebloune Y, & Moussaoui A. Molecular Characterization of *Aspergillus flavus* Strains Isolated from Animal Feeds. Pol. J. Microbiol. 2022;71(4): 589–599. 10.33073/pjm-2022-048

5. Rudramurthy SM, Paul RA, Chakrabarti A, Mouton JW, & Meis JF. Invasive Aspergillosis by *Aspergillus flavus*: Epidemiology, Diagnosis, Antifungal Resistance, and Management. J. Fungi. 2019;5(3): 55. 10.3390/jof5030055

6. Pickova D, Ostry V, & Malir F. A Recent Overview of Producers and Important Dietary Sources of Aflatoxins. Tox. 2021;13(3): 186. 10.3390/toxins13030186

7. Cary JW, Gilbert MK, Lebar MD, Majumdar R, & Calvo AM. *Aspergillus flavus* Secondary Metabolites: More than Just Aflatoxins. J. Food Saf. 2018;6(1): 7–32. 10.14252/foodsafetyfscj.2017024

8. Ribas D, Piérri Spolti Del EM, Donato KZ, Schrekker HS, & Fuentefria AM. Is the emergence of fungal resistance to medical triazoles related to their use in the agroecosystems? A mini review. Braz. J. Microbiol. 2016;47(4): 793–799. 10.1016/j.bjm.2016.06.006

9. Choi MJ, Won EJ, Joo MY, Park YJ, Kim SH, Shin MG, et al. Microsatellite Typing and Resistance Mechanism Analysis of Voriconazole-Resistant *Aspergillus flavus* Isolates in South Korean Hospitals. Antimicrob. Agents Chemother. 2019;63(2): e01610–18. 10.1128/AAC.01610-18

10. Toda M, Beer KD, Kuivila KM, Chiller TM, & Jackson BR. Trends in Agricultural Triazole Fungicide Use in the United States, 1992–2016 and Possible Implications for Antifungal-Resistant Fungi in Human Disease. Environ. Health Perspect. 2021;129(5). 10.1289/ehp7484

11. Fountain JC, Clevenger J, Nadon B, Youngblood RC, Korani W, Chang PK., et al. Two New *Aspergillus flavus* Reference Genomes Reveal a Large Insertion Potentially Contributing to Isolate Stress Tolerance and Aflatoxin Production. G3 (Bethesda, Md.). 2020;10(10): 3515–3531. 10.1534/g3.120.401405

12. Gangurde SS, Korani W, Bajaj P, Wang H, Fountain JC, Agarwal G, et al. *Aspergillus flavus* pangenome (AflaPan) uncovers novel aflatoxin and secondary metabolite associated gene clusters. BMC Plant Biol. 2024;24(1). 10.1186/s12870-024-04950-8

13. Mitema A, & Aliye Feto N. Molecular and Vegetative Compatibility Groups Characterization of *Aspergillus flavus* Isolates from Kenya. AIMS Microbiol. 2020;6(3): 231–250. 10.3934/microbiol.2020015

14. Hatmaker EA, Barber AE, Drott MT, Sauters TJC, Gumilang A, Alastruey-Izquierdo A, Garcia-Hermoso D, et al. Population structure in a fungal human pathogen is potentially linked to pathogenicity. Nat. Comm. 2025;16(1). 10.1038/s41467-025-62777-9

15. Horn BW, Gell RM, Singh R, Sorensen RB, & Carbone I. Sexual Reproduction in *Aspergillus flavus* Sclerotia: Acquisition of Novel Alleles from Soil Populations and Uniparental Mitochondrial Inheritance. PLOS ONE. 2017;11(1): e0146169–e0146169. 10.1371/journal.pone.0146169

16. Slatkin M. Linkage disequilibrium — understanding the evolutionary past and mapping the medical future. Nat. Rev. Genet. 2008;9(6): 477–485. 10.1038/nrg2361

17. Mvogo Nyebe RA, Kumar A, Ngonkeu Mangaptche EL, Kumar S, Velmurugan S, Krishnappa C, et al. Multi-omics characterization of aflatoxigenic *Aspergillus* from grains and rhizosphere of maize across agroecological zones of Cameroon. Sci. Rep. 2025;15(1): 18407. 10.1038/s41598-025-97296-6

18. Ntsoli PG, Boat Bedine MA, Baleba CC, Tchatcho Ngalle SF, Djoko Kouam I, Titti RW, et al. Postharvest Practices, Perceptions, and Knowledge of Mycotoxins among Groundnut Farmers in the Adamawa, Centre, and North Regions of Cameroon. Scientifica. 2024;2024: 1–16. 10.1155/2024/5596036

19. Pouokam G, Lemnyuy Album W, Ndikontar A, & Sidatt M. A Pilot Study in Cameroon to Understand Safe Uses of Pesticides in Agriculture, Risk Factors for Farmers’ Exposure and Management of Accidental Cases. Tox. 2017;5(4): 30. 10.3390/toxics5040030

20. Mianrood IB, Gharebagh FJ, Khodavaisy S, Ahmadian M, & Darazam IA. Meta-analysis of antifungal resistance patterns of *Aspergillus* species in Iran. J. Infect. Public Health. 2025;18(9): 102838. 10.1016/j.jiph.2025.102838

21. Atehnkeng J, Donner M, Ojiambo PS, Ikotun B, Augusto J, Cotty PJ, et al. Environmental distribution and genetic diversity of vegetative compatibility groups determine biocontrol strategies to mitigate aflatoxin contamination of maize by *Aspergillus flavus*. Microb. Biotechnol., 2015;9(1): 75–88. 10.1111/1751-7915.12324

22. Fantong W, Fouepe A, ISSA N first name, Djomou S, Banseka H, An K, et al. Temporal pollution by nitrate (NO 3), and discharge of springs in shallow crystalline aquifers: Case of Akok Ndoue catchment, Yaounde (Cameroon). Afr. J. Environ. Sci. Technol. 2013;7:175–91. doi:10.5897/AJEST2013.1421

23. Xiao W, Gong D ying, Mao B, Du X miao, Cai LL, Wang M yu, et al. Sputum signatures for invasive pulmonary aspergillosis in patients with underlying respiratory diseases (SPARED): study protocol for a prospective diagnostic trial. BMC Infect Dis. 2018;18:271. doi:10.1186/s12879-018-3180-z PubMed PMID: 29890956; PubMed Central PMCID: PMC5996557.

24. Vergidis P, Moore C, Rautemaa-Richardson R, Richardson M. High-volume Sputum Culture for the Diagnosis of Pulmonary Aspergillosis. Open Forum Infect Dis. 2017;4(Suppl 1):S609. doi:10.1093/ofid/ofx163.1598 PubMed PMID: null; PubMed Central PMCID: PMC5631283.

25. Ashu EE, Korfanty GA, Xu J. Evidence of unique genetic diversity in *Aspergillus fumigatus* isolates from Cameroon. Mycoses. 2017;60(11):739–48. doi:10.1111/myc.12655

26. Samarasinghe H, Korfanty G, Xu J. Isolation of Culturable Yeasts and Molds from Soils to Investigate Fungal Population Structure. JoVE. 2022;(183):63396. doi:10.3791/63396

27. Temesgen T, Chala B. Isolation and Identification of Fungi from Peanut Samples Sold at Adama City Markets, Oromia Regional State, Ethiopia. Int. J. Sci. Res. Biol. Sci. 2020;7(3):79–84. https://ijsrbs.isroset.org/index.php/j/article/view/372

28. Nguyen XH, Nguyen TMN, Nguyen DH, Nguyen QC, Cao TT, Pham TTH, et al. Identification and characterization of *Aspergillus niger* causing collar rot of groundnut (*Arachis hypogaea*). Biodiversitas Journal of Biological Diversity. 2023;24(5):5. doi:10.13057/biodiv/d240507

29. Of I, Ng O, Fi O, Ae A. Fungal Contaminants Associated with Groundnut (Arachis Hypogaea) Seeds. J Bioinform Syst Biol. 2021;04(04). doi:10.26502/jbsb.5107029

30. Doilom M, Guo JW, Phookamsak R, Mortimer PE, Karunarathna SC, Dong W, et al. Screening of Phosphate-Solubilizing Fungi From Air and Soil in Yunnan, China: Four Novel Species in *Aspergillus, Gongronella, Penicillium*, and *Talaromyces*. Front Microbiol. 2020;11. doi:10.3389/fmicb.2020.585215

31. Xu J, Ramos AR, Vilgalys R, & Mitchell TG. Clonal and spontaneous origins of fluconazole resistance in *Candida albicans*. J. Clin. Microbiol. 2000;38(3): 1214–1220. 10.1128/JCM.38.3.1214-1220.2000

32. Halvaeezadeh M, Jalaee GA, Fatahinia M, & Zarei Mahmoudabadi A. Microsatellite Genotyping of Clinical and Environmental *Aspergillus flavus*. Jundishapur J. Microbiol. 2025;19(1). 10.5812/jjm-167602

33. Fan Y, Korfanty GA, & Xu J. Genetic Analyses of Amphotericin B Susceptibility in *Aspergillus fumigatus*. J. Fungi. 2021;7(10): 860–860. 10.3390/jof7100860

34. Jørgensen KM, Helleberg M, Hare RK, Jørgensen LN, & Arendrup MC. Dissection of the Activity of Agricultural Fungicides against Clinical *Aspergillus* Isolates with and without Environmentally and Medically Induced Azole Resistance. J. Fungi. 2021;7(3): 205. 10.3390/jof7030205

35. Goor RM, Hoffman D, & Riley GR. Novel Method for Accurately Assessing Pull-up Artifacts in STR Analysis. Forensic Sci. Int. Genet. 2021;51: 102410. 10.1016/j.fsigen.2020.102410

36. Peakall R, & Smouse P. GenAlEx 6.5: genetic analysis in Ex-cel. Population genetic software for teaching and research-an up-date. Bioinform. 2012;28(19): 2537–2539. doi:10.1093/bioinformatics/bts460.

37. Kamvar ZN, Tabima JF, & Grünwald NJ. Poppr: an R package for genetic analysis of populations with clonal, partially clonal, and/or sexual reproduction. PeerJ. 2014;2: e281. doi:10.7717/peerj.281.

38. Letunic I, & Bork P. Interactive Tree of Life (iTOL) v5: an online tool for phylogenetic tree display and annotation. Nucleic Acids Res. 2021;49(W1): 293–296. doi:10.1093/nar/gkab301.

39. Agapow PM, & Burt A. Indices of multilocus linkage disequilibrium. Mol. Ecol. Notes. 2001;1(1–2): 101–102. doi:10.1046/j.1471-8278.2000.00014.x.

40. Duong TMN, Nguyen PT, Le TV, Nguyen HLP, Nguyen BNT, Nguyen BPT, et al. Drug-Resistant *Aspergillus flavus* Is Highly Prevalent in the Environment of Vietnam: A New Challenge for the Management of Aspergillosis? J. Fungi. 2020;6(4): 296. 10.3390/jof6040296

41. Meireles LM, Soares FS, de A, & Scherer R. Exposure to Tebuconazole Drives Cross-Resistance to Clinical Triazoles in *Aspergillus fumigatus*. ACS Omega. 2025;10(42): 50407– 50414. 10.1021/acsomega.5c07812

42. Drott MT, Rush TA, Satterlee TR, Giannone RJ, Abraham PE, Greco C, et al. Microevolution in the pansecondary metabolome of *Aspergillus flavus* and its potential macroevolutionary implications for filamentous fungi. Proc. Natl. Acad. Sci. U. S. A. 2021;118(21). 10.1073/pnas.2021683118

43. Blacket MJ, Robin C, Good RT, Lee SF, & Miller AD. Universal primers for fluorescent labelling of PCR fragments-an efficient and cost-effective approach to genotyping by fluorescence. Mol. Ecol. Resour. 2012;12(3): 456–463. 10.1111/j.1755-0998.2011.03104.x

44. Wang Y, Han L, Gong J, Liu L, Miao BB, Xu J. Research advances and public health strategies in China on WHO priority fungal pathogens. Mycology. 2025;16(4): 1437–1477. 10.1080/21501203.2025.2561612

